## Supplementary Information for "Deep model predictive control of gene expression in thousands of single cells"

**Supplementary Figures**

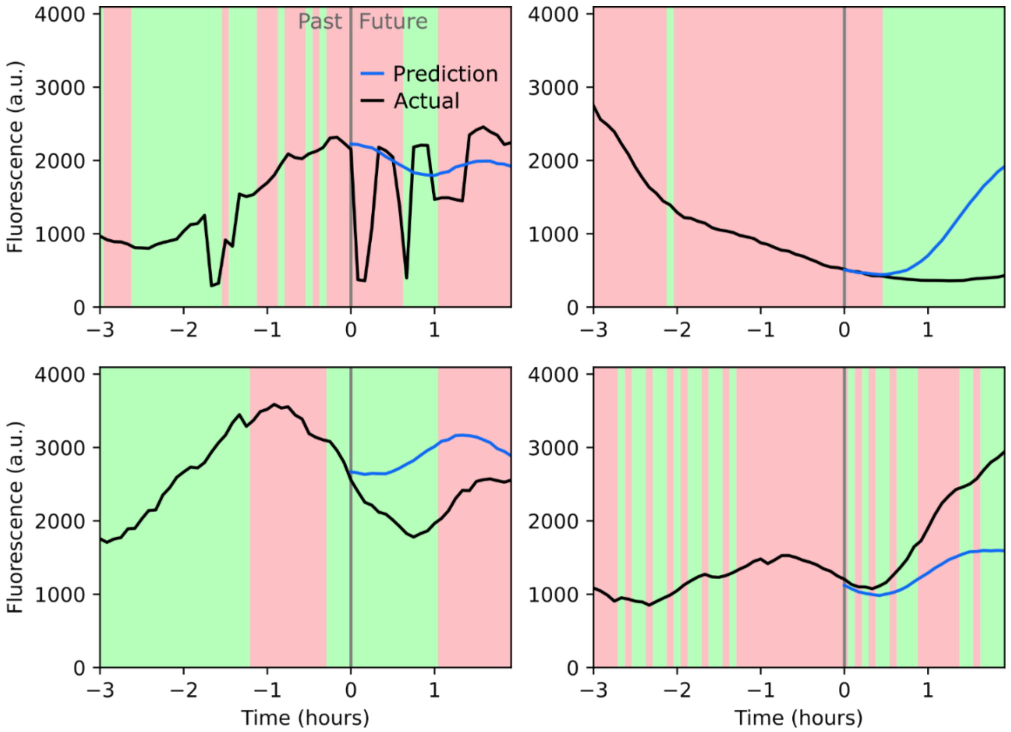


**Figure S1**. Predictions for the 95th root mean square error percentile of the 2-hour horizon model. Four representative examples are shown: In the top left panel, image analysis errors result in unpredictable jumps in fluorescence levels, and in the other 3 panels the cell "responsiveness" seems to suddenly change compared to past behavior. Red/green background colors represent optogenetic stimulations.

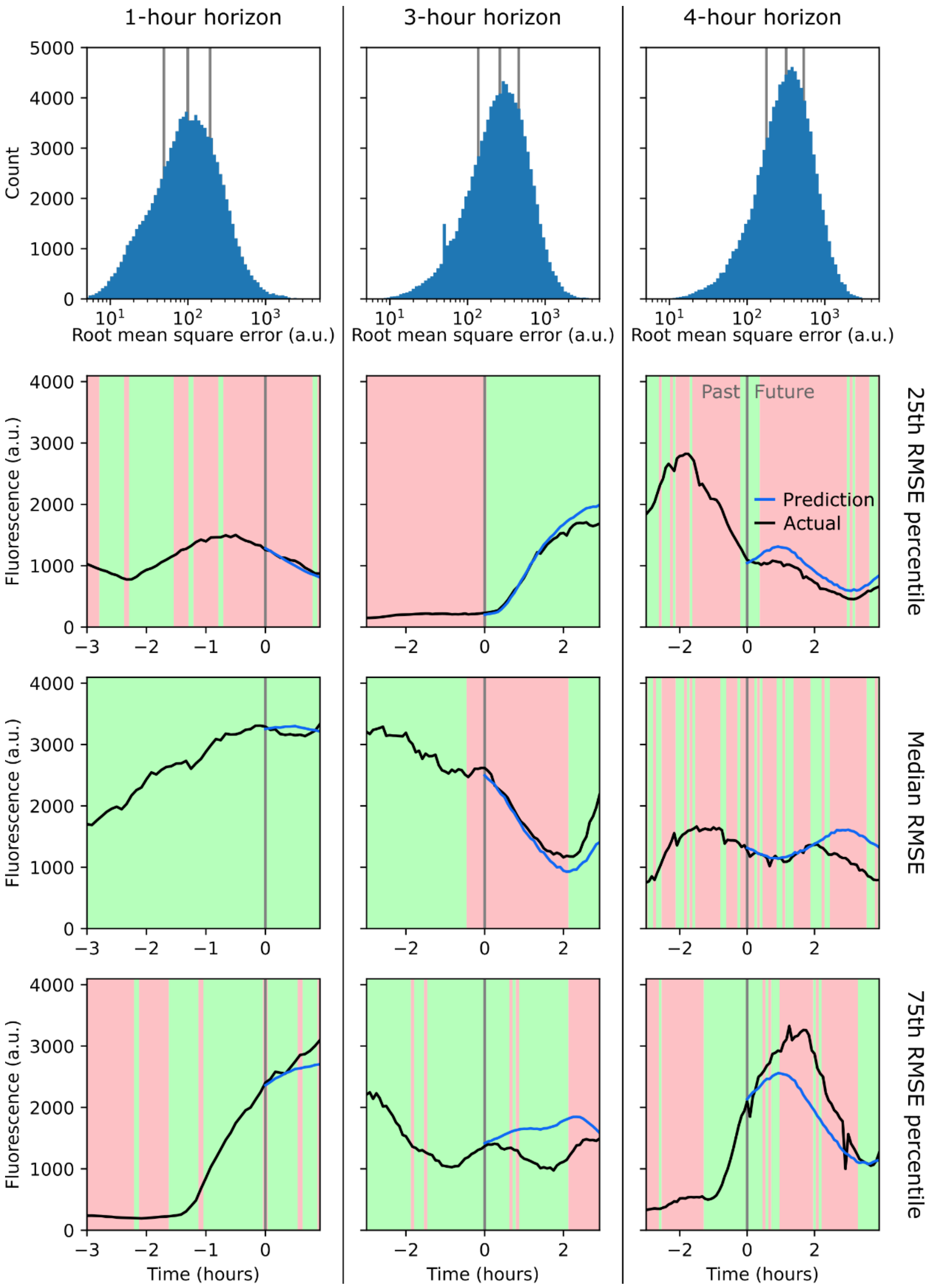


**Figure S2**. Performance of 1, 3, and 4-hour horizon models. Top row is the histogram of the root mean square error between 100,000 model predictions on the validation dataset and their ground truth. Vertical gray lines represent the 25th, median, and 75th percentile of the root mean square error, from left to right. Below each histogram are illustrative predictions at the 25th, median, and 75th percentile of the error for each of the different prediction horizon models.

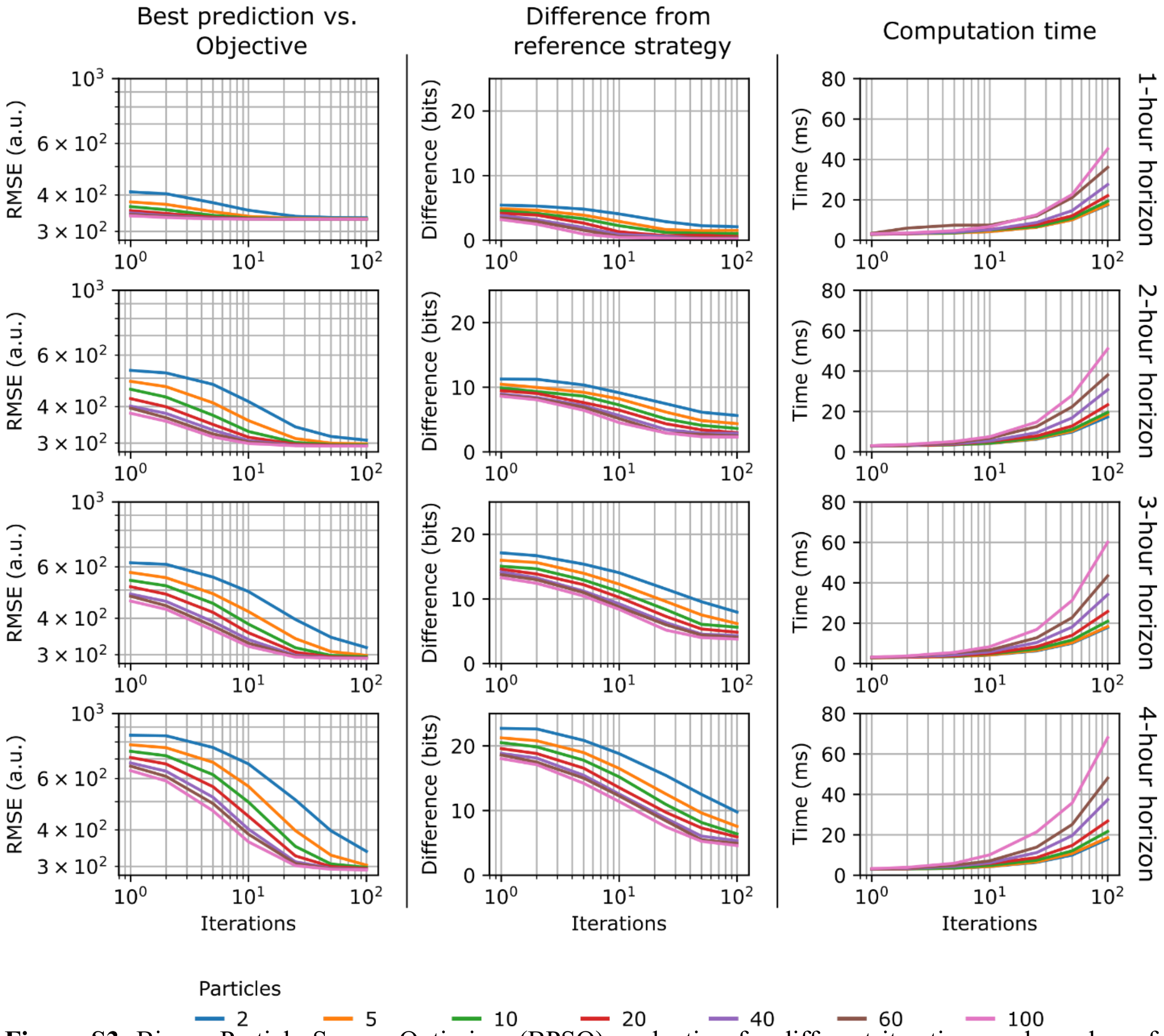


**Figure S3**. Binary Particle Swarm Optimizer (BPSO) evaluation for different iterations and number of particles. All evaluations were run on 524 single-cell control samples derived from experimental data. Briefly, Particle Swarm Optimizers evaluate a number of potential solutions in parallel (particles, colored lines) over a certain number of iterations (x-axis) and finally select the best solution. The "best" solution selected by the BPSO is whichever optogenetic stimulations strategy is predicted to drive fluorescence closest to the objective. Each row from top to bottom is for the 1, 2, 3, and 4-hour horizon model. Left column represents the average root mean square error between the prediction from the "best" strategy as determined by the BPSO and the control objective. Central column shows the average difference between the "best" strategy determined by the BPSO and a reference strategy (e.g. a sequence of [red, green, red, green] and a sequence of [red, red, red, green] stimulations would differ by 1 bit). The "best" reference strategy was determined by running the BPSO with 500 particles and 200 iterations. This represents an extreme case with many particles and a large number of iterations to identify a near-optimal strategy. The right column shows the execution time of the BPSO for a single control sample. Computation times are faster than during our experiments because here all 524 single-cell samples are evaluated in parallel, while in practice, data are processed for 27-28 cells at a time since that corresponds to the number of mother machine chambers in a single field of view. We concluded that using 40 particles over 25 iterations was optimal for our applications and across horizons.

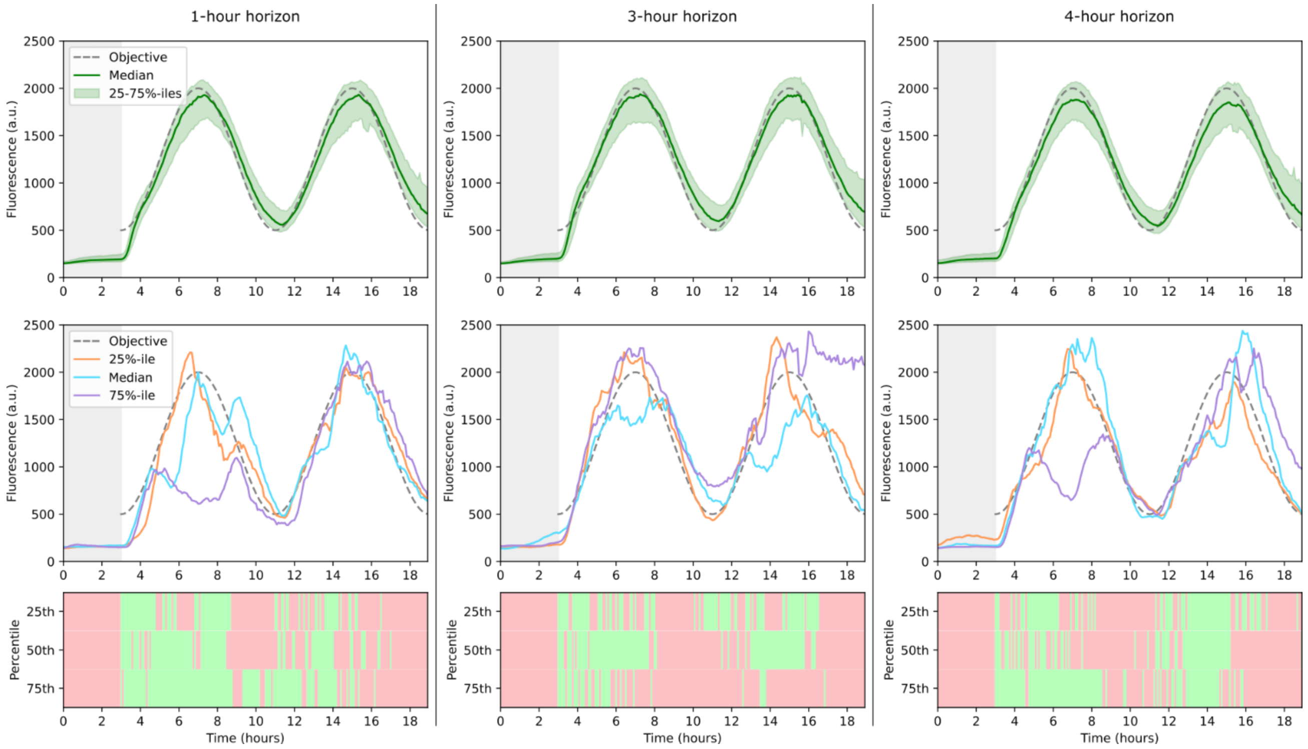


**Figure S4**. Control performance for different horizon controllers. 1, 3, and 4-hour horizons are shown from left to right (n = 524, 524, and 496 cells). The top row is the control performance at the population level. The gray dashed curve represents the control objective. The solid green curve represents the median fluorescence of the population. The shaded area represents 25$^{th}$ to 75$^{th}$ percentiles of the population fluorescence. The second row is the control performance at the single-cell level. Colored solid curves show representative single-cell fluorescence trajectories in the 25$^{th}$, median, and 75$^{th}$ percentiles of control accuracy. Red and green background colors in the third row represent the optogenetic stimulation sequences that were applied by the controller for the representative three single-cell trajectories, with the 75$^{th}$ percentile, median, and 25$^{th}$ percentile shown from top to bottom.

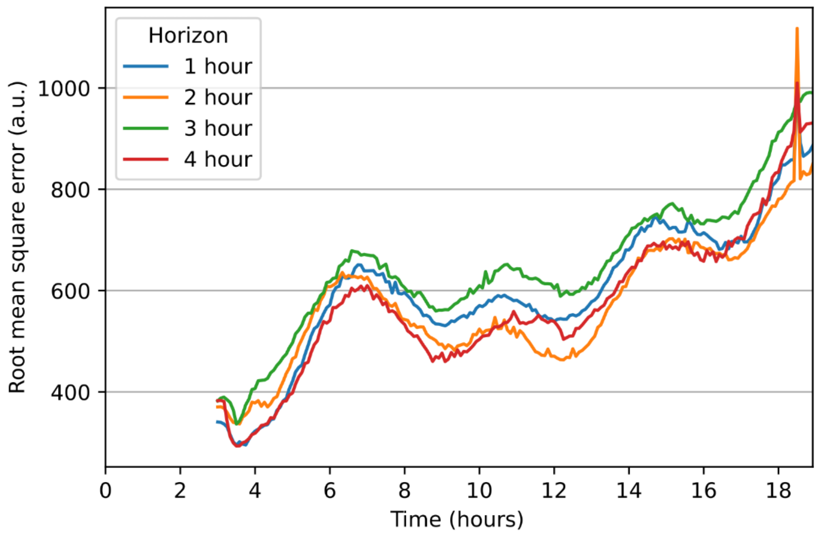


**Figure S5**. Root mean square error over time between cell fluorescence and single sinewave objective per timepoint, for controllers with 1, 2, 3 and 4 hour horizons.

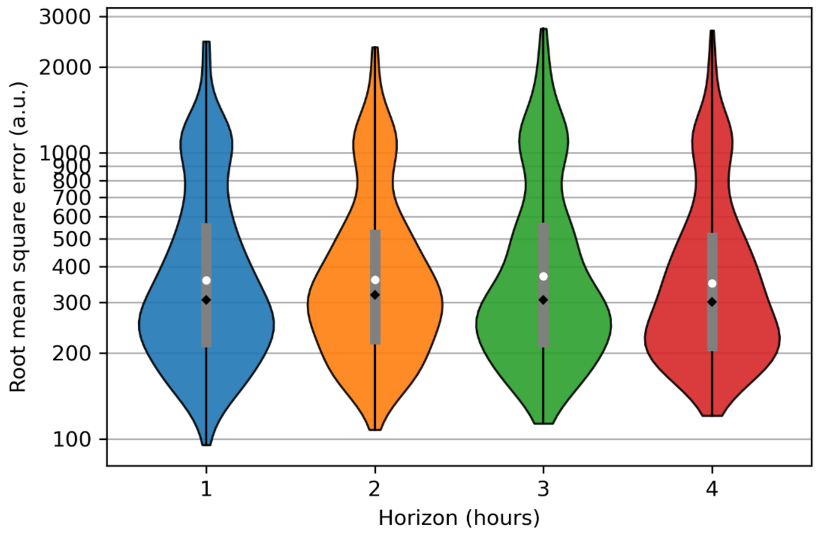


**Figure S6** – Violin plot of distribution of root mean square error between single cell fluorescence and sinewave objective, for controllers with 1, 2, 3, and 4 hour horizons.

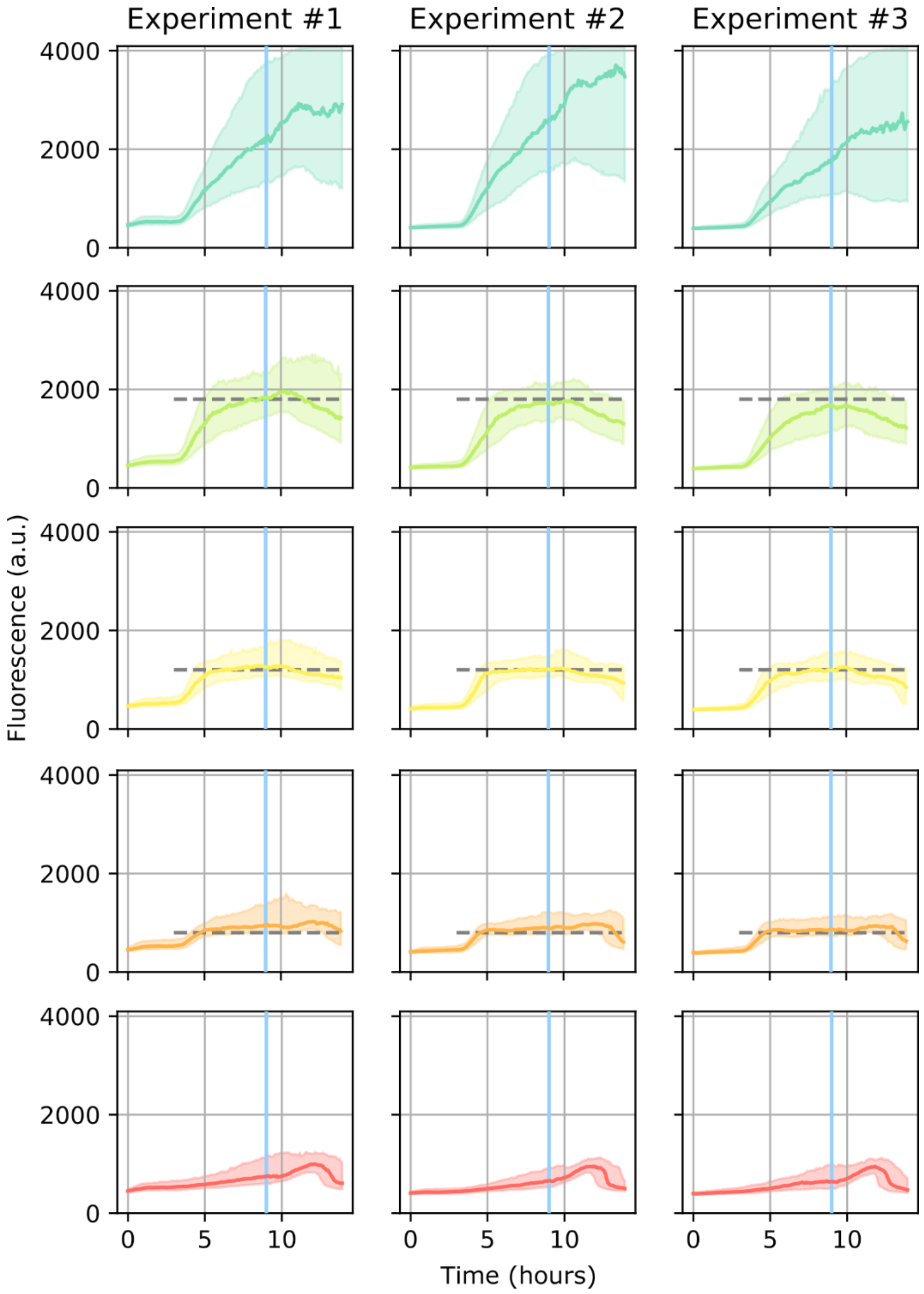


**Figure S7**. Population response with *tetA-gfp* control per category and replicate experiment. Top and bottom row represent the cells subjected to constant green and constant red optogenetic stimulations. The middle rows represent cells controlled at 1800, 1200, and 800 units of fluorescence. All cells received exclusively red stimulations for equilibration until t = 3h. Horizontal dashed gray lines represent control objectives. Vertical blue lines represent the time of tetracycline addition at t = 9h. Solid colored curves represent the population median fluorescence. The shaded areas represent the 25$^{th}$ to 75$^{th}$ fluorescence percentiles.

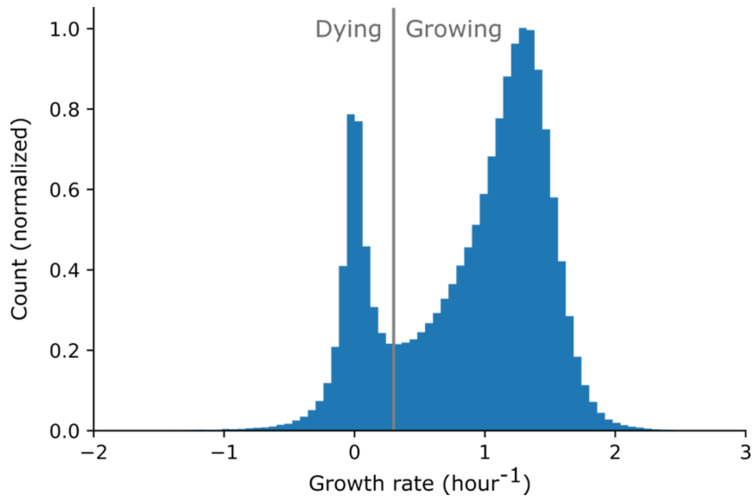


**Figure S8**. Histogram of growth rates for *tetA-gfp* control experiments, across all replicates, cell populations, and timepoints. Single cell growth rates were smoothed with a median filter sliding over a one-hour time window. The vertical gray line represents the 0.3 hour$^{-1}$ "growing" vs. "dying" threshold.

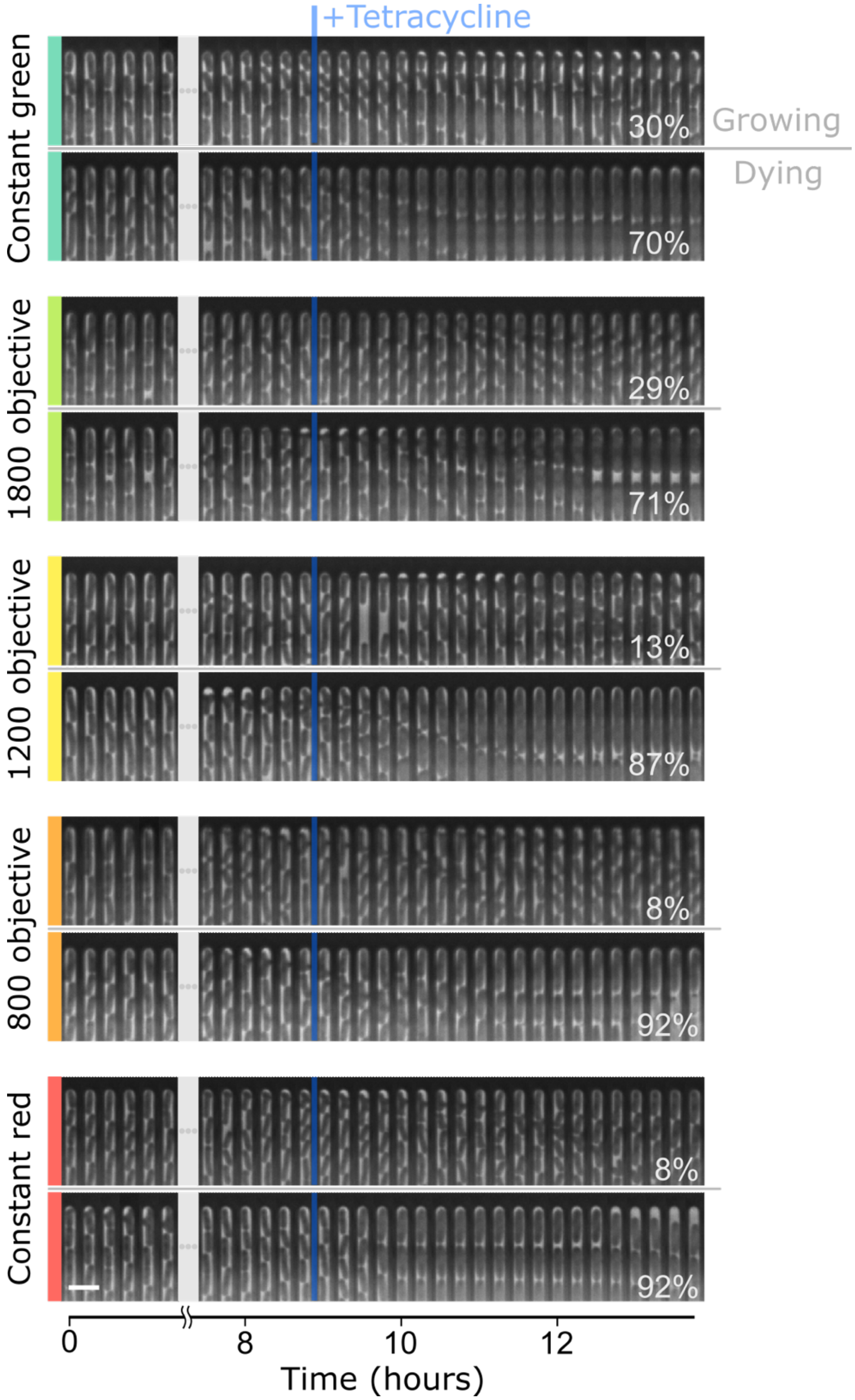


**Figure S9**. Representative kymographs of cells in the “growing” or “dying” sub-populations for each category of cells. Listed values indicate percentage of cells in each sub-population at the end of the experiment. Scale bar, 5 µm.

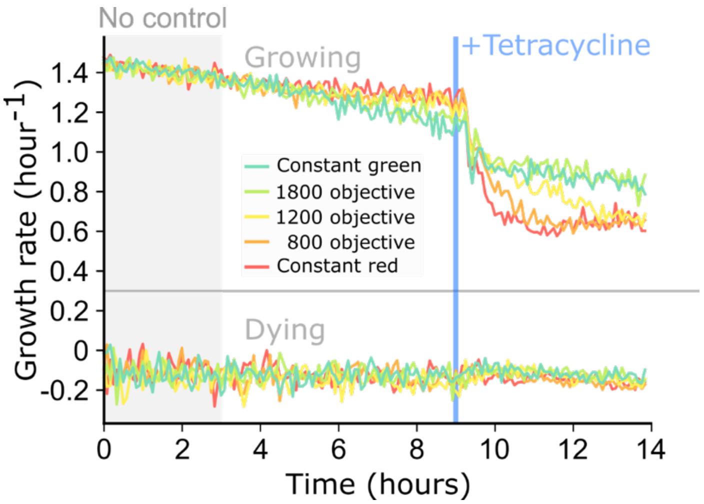


**Figure S10**. Median growth rates over time for cells in the "growing" or the "dying" sub-populations. Data from all replicates were merged. The growth rate over time was not smoothed.

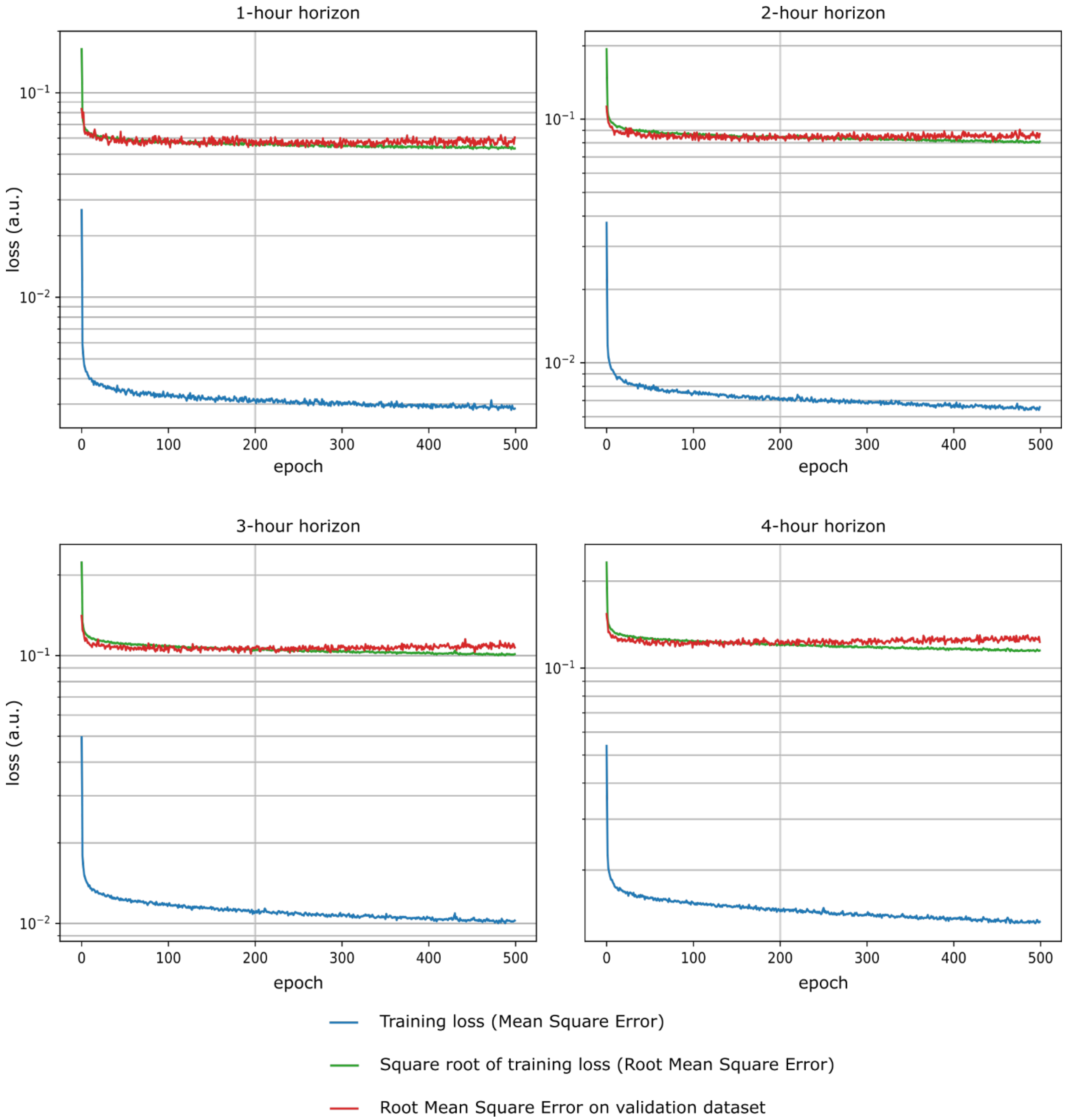


**Figure S11**. Training loss and validation error of the 1, 2, 3, and 4-hour horizon models. The models were trained for up to 500 epochs. At the end of each epoch, the model was evaluated against 10,000 samples from the validation dataset. The model with the best root mean square error against validation data over time was saved to disk, and that model was used for timeseries forecasting evaluation. For all horizons, we observed that around 200 epochs the validation error seemed to plateau or even increase while the training loss kept decreasing.

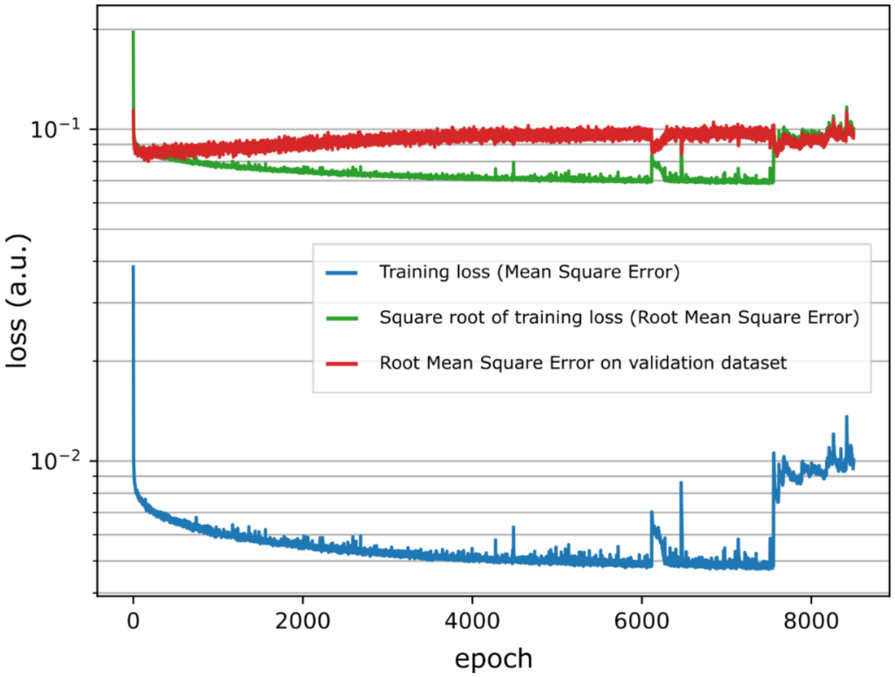


**Figure S12**. Training loss and validation error of the 2-hour horizon model up to 8,504 epochs. The maximum number of epochs was set to 10,000, with a patience parameter that stopped the training 1,000 epochs after the training loss stopped decreasing. Validation error increases as loss decreases, indicating model overfitting.

**Supplementary Text**

Random optogenetic stimulation sequences for training set experiments.

For our training sets, we want the pre-determined optogenetic stimulations sequences to be as diversified as possible, but also for the sequences to often contain relatively long sub-sequences with only red or green stimulations. Simply generating a Bernoulli sequence of random coin flips, even biased, would rapidly alternate between red and green stimulations for most of the sequence. Instead, we used a bounded one-dimensional random walk to generate our training stimulation sequences:

1. For each cell, a uniformly random sequence of either +1 or -1 values was generated. The sequence was generated for up to 36 hours of stimulations every 5 minutes, or 432 total values, although we only ended up running training experiments for up to 24 hours. An example sequence would be:

   -1, -1, 1, -1, -1, -1, -1, -1, 1, 1, -1, 1, -1, -1, -1, 1, 1, 1, 1, 1, -1, 1, -1, 1, -1, -1, 1, 1, -1, …

2. This random sequence was then cumulatively summed over time, however that sum was bounded to maximum and minimum values, respectively positive and negative. This was meant to prevent cumulatively summed values from growing too large or too small. Different sets of maximum and minimum values between +4 and -3 were used randomly for different cells, to generate different distributions of red and green stimulations. The cumulative sum of the sequence above bounded to [-2, 2] would be:

   0, -1, -2, -1, -2, -2, -2, -2, -2, -1, 0, -1, 0, -1, -2, -2, -1, 0, 1, 2, 2, 1, 2, 1, 2, 1, 0, 1, 2, …

3. Finally, the values are binarized. All values greater than or equal to 0 are set as green stimulations while negative values are set to red stimulations. The final sequence for the example would be: G, R, R, R, R, R, R, R, R, R, G, R, G, R, R, R, R, G, G, G, G, G, G, G, G, G, G, G, G, G, …

At the beginning of each sequence we appended 3 hours (=36 time points) of only red stimulations to mimic control conditions and make sure cells started in a repressed state.

Single-cell feature normalization for neural network processing

Input normalization is good practice for training and using neural networks. We used the following formulas to ensure single-cell timeseries data were consistently normalized to the same range of values:

- **Fluorescence**: Single-cell fluorescence measurements were limited to [0, 4095], the dynamic range of the data. Fluorescence features values (mother cell fluorescence, average chamber fluorescence, and chamber fluorescence standard deviation) were thus simply normalized linearly:

  $$x_{norm} = x_{raw}/4095$$

  Where $x_{raw}$ is the initial measured value for a given feature and time point, and $x_{norm}$ is the resulting normalized value that is used as input for the neural network, both for training and on-the-fly for feedback control.

- **Cell area:** Mother cell area, measured in pixels, theoretically has no clear upper bound. After analyzing area distributions, we used a negative exponential function:

$$x_{norm} = 1 - 10^{-x_{raw}/3000}$$

- **Chamber cell count**: The number of cells in the chamber at any given timepoint also theoretically has no clear upper bound. We also used a negative exponential function with a different normalization factor:

$$x_{norm} = 1 - 10^{-x_{raw}/9}$$

- **Chamber image sharpness**: Image sharpness, measured as the average of the Laplacian of the image, has no clear upper or lower bound. We used the following formula to normalize sharpness to [0, 1]:

$$x_{norm} = \frac{1}{1 + e^{-x_{raw}+3.6}}$$

**Supplementary Movie Captions**

**Movie S1**. Control strategies over time. All three cells from Fig. 3C are represented, with trajectories in the 25$^{th}$ percentile, median, and 75$^{th}$ percentile of control accuracy shown from top to bottom. Dashed gray curves show the control objective. Sliding vertical gray lines represent the current time point. Red and green background colors indicate the optogenetic stimulation sequences that were applied before the gray line, and the evolving control strategy after the gray line. Colored solid curves show measured single-cell fluorescence before the gray line, and the predicted response to the control strategy after the gray line.

**Movie S2**. Expanding concentric sinewaves population patterning. The left panel shows the control objectives that were assigned to each cell. The center panel shows measured fluorescence values for single cells subjected to deep model predictive control in the experiment (n = 10,000 cells). The two right panels show fluorescence and optogenetic stimulations for two representative controlled cells at pixel coordinates (15, 15) and (75, 50). The movies are mirrored vertically with respect to Fig. 3D to prevent the lines indicating representative cells from obscuring the movie.

**Movie S3**. *2001: A Space Odyssey* scene reproduction. The left panel shows the control objectives that were assigned to each cell. The right panel shows measured fluorescence values for single cells subjected to deep model predictive control in the experiment (n = 10,000 cells).

**Movie S4**. Growth rate violin plots for all five control categories before and after tetracycline addition at t = 9h. Horizontal gray line represents the 0.3 $h^{-1}$ "growing" vs. "dying" growth rate threshold. For visual clarity, the top and bottom 0.25% of the growth rate distributions were filtered out.
